# SARS-CoV-2 infection paralyzes cytotoxic and metabolic functions of the immune cells

**DOI:** 10.1101/2020.09.04.282780

**Authors:** Yogesh Singh, Christoph Trautwein, Rolf Fendel, Naomi Krickeberg, Georgy Berezhnoy, Rosi Bissinger, Stephan Ossowski, Madhuri S. Salker, Nicolas Casadei, Olaf Riess, the Deutsche COVID-19 OMICS Initiative (DeCOI)

## Abstract

The SARS-CoV-2 virus is the causative agent of the global COVID-19 infectious disease outbreak, which can lead to acute respiratory distress syndrome (ARDS). However, it is still unclear how the virus interferes with immune cell and metabolic functions in the human body. In this study, we investigated the immune response in acute or convalescent COVID19 patients. We characterized the peripheral blood mononuclear cells (PBMCs) using flow cytometry and found that CD8^+^ T cells were significantly subsided in moderate COVID-19 and convalescent patients. Furthermore, characterization of CD8^+^ T cells suggested that patients with a mild and moderate course of the COVID-19 disease and convalescent patients have significantly diminished expression of both perforin and granzyme A in CD8^+^ T cells. Using ^1^H-NMR spectroscopy, we characterized the metabolic status of their autologous PBMCs. We found that fructose, lactate and taurine levels were elevated in infected (mild and moderate) patients compared with control and convalescent patients. Glucose, glutamate, formate and acetate levels were attenuated in COVID-19 (mild and moderate) patients. In summary, our report suggests that SARS-CoV-2 infection leads to disrupted CD8^+^ T cytotoxic functions and changes the overall metabolic functions of immune cells.

## Introduction

The first cases of severe acute respiratory coronavirus-2 (SARS-CoV-2) infection appeared in December 2019, in Wuhan, China^1^. This zoonotic virus has infected by now more than 127.8 million people (30.03.2021) and has resulted more than 2.78 million^2,3^ death worldwide. The containment of the pandemic is challenging and is still growing with roughly 200,000 or more new infections being reported daily since July 2020^2,3^. There is an urgent need for a better understanding of immunopathology, as SARS-CoV-2 has become the leading cause of morbidity (long COVID syndrome) and mortality in many countries.

Coronaviruses (CoV) are a large family of viruses that can cause illnesses such as the common cold and seasonal influenza^4^. Pathologically, SARS-CoV-2 typically infects via angiotensin-converting enzyme 2 (ACE2)-expressing nasal epithelial cells in the upper respiratory tract and type II alveolar epithelial cells in patients exhibiting pneumonitis^1,5^. The most severe disease courses led frequently to death but, not exclusively in older patients with and without risk conditions. The primary symptoms of SARS-CoV-2 infections are fatigue, fever, sore throat, dry cough, loss of smell and taste within 5-21 days of incubation of the virus^6-9^. COVID-19 symptoms are heterogeneous and range from asymptomatic to mild, moderate, and severe pathological symptoms, presenting with or without pneumonia^10,11^. However, most infected people develop mild to moderate illness and recover without hospitalization^12,13^. High serum levels of IL-6, IL-8, IL-10, TNF-α cytokines and an immune hyper-responsiveness referred to as a ‘cytokine storm’ is connected with poor clinical outcome^14,15^. Predominantly, older COVID-19 patients can develop acute severe respiratory distress syndrome (ARDS) due to a cytokine storm which is a life-threatening situation, requiring ventilation and intensive care support^16-20^.

Several break-through discoveries have extended our understanding as to how the virus takes advantage of the host and modulates immunity^12,19,21-25^. Recovered COVID-19 patients have an increased number of antibody-secreting cells and activated CD4^+^ and CD8^+^ T cells. Further, Immunoglobulin M (IgM) and SARS-CoV-2 reactive IgG antibodies were also detected in blood before full symptomatic recovery^26-28^. Most severely affected COVID-19 patients had a lower T cell but elevated B cell counts when compared with healthy controls^13,14,29,30^. Interestingly, patients with mild symptoms were also shown to have increased T and B cells compared with severely affected patients^26,29-31^. There could be several reasons for different disease outcomes including an over-activated innate or hyper-activated adaptive immune response leading to cytokine storms and resulting in severe injury to the lungs^10,13,25,32^. Despite several ongoing efforts, the immunological mechanisms of the host-pathogen interaction are not well understood^33^.

There is an intricate balance between the metabolic state of immune cells and generation of a robust immune response^19,34-37^. CD8^+^ T cells require energy to proliferate and accomplish their effective functions^38^. Most propagating cells such as lymphocytes utilize the most abundant energy substrates including, glucose, lipids, and amino acids^39^. In response to SARS-CoV-2 and other virus infections, CD8^+^ T cells play a pivotal role in profound growth and proliferation to generate their effective functional cells which can produce copious amounts of effector molecules such as cytokines and cytotoxic granules^30,38-40^. An activated immune system is coupled with a change in metabolic reprogramming to produce enough energy needed during (viral) infection^38,39^. Proliferating T cells ferment glucose to lactate even in the presence of oxygen to meet high energy demands^34,37-39^. Furthermore, glucose and glutamine are involved in the hexosamine biosynthetic pathway, which regulates the production of uridine diphosphate N-acetyl glucosamine necessary for T cell clonal expansion and function^41^. The synthesis of lactate intracellularly is crucial for T cells to have an increased glycolytic flux^38^.

Peripheral blood mononuclear cells (PBMCs) can be analyzed to measure the physiological dysfunctionality of an individual and can serve as a biomarker^42^. Consequently, the metabolic status of lymphocytes could help to predict disease severity or to select the optimal therapeutic intervention to boost the immune function during infection. Generally, most of the metabolism-related functions in PBMCs during SARS-CoV-2 infections were inferred based on transcriptomics analysis^34,43^ and no functional data (biochemical level) have been presented. Therefore, understanding the kinetics of the adaptive immune response as well as the metabolic functions during SARS-CoV-2 infection will help to elucidate the host immune response to SARS-CoV-2 infection. In this study, using flow cytometry and proton nuclear magnetic resonance (^1^H-NMR) spectroscopy, we characterized the PBMCs from SARS-CoV-2 infected and convalescent patients for their immunophenotypic and metabolic functions.

## Results

### Characteristics of study participants

PBMCs were isolated and cryopreserved from blood samples obtained from COVID-19 patients suffering from mild (‘Mild (outpatient)’) or moderate/severe (‘Moderate (inpatient)’) disease or were already recovered (‘Convalescent’) and from healthy controls (‘HC’). Classification of disease severity for this analysis was based on the requirement of hospitalization. Patients with mild COVID-19 were recruited within three days after confirmation of infection by RT-qPCR. From moderate to severe COVID-19 patient blood samples were collected one week after their hospital admittance. The moderate patients were admitted to the hospital requiring medical care; however, they did not need ventilation or O2 supply. Recovered patients were included based on a positive SARS-CoV-2 antibody testing. Study participant characteristics are described in Table 1.

**Table 1:**
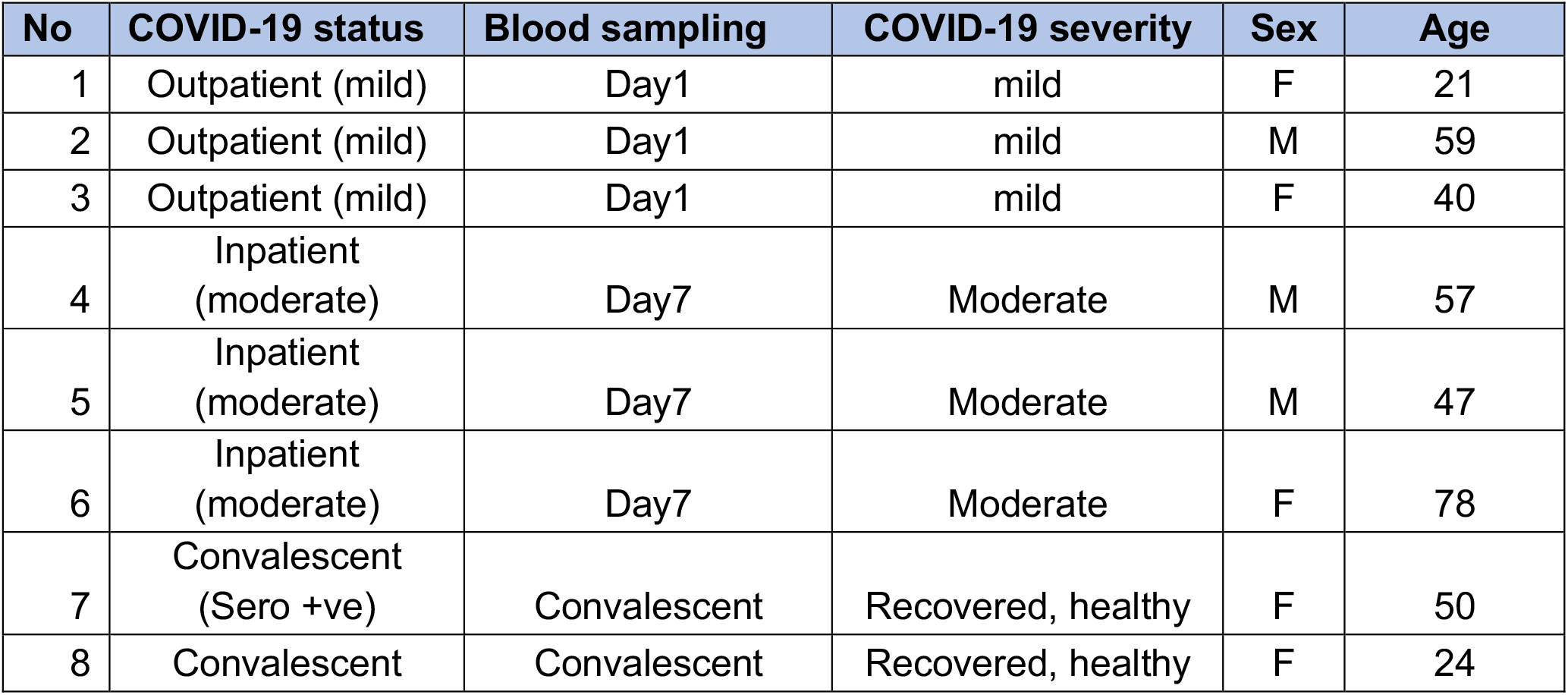
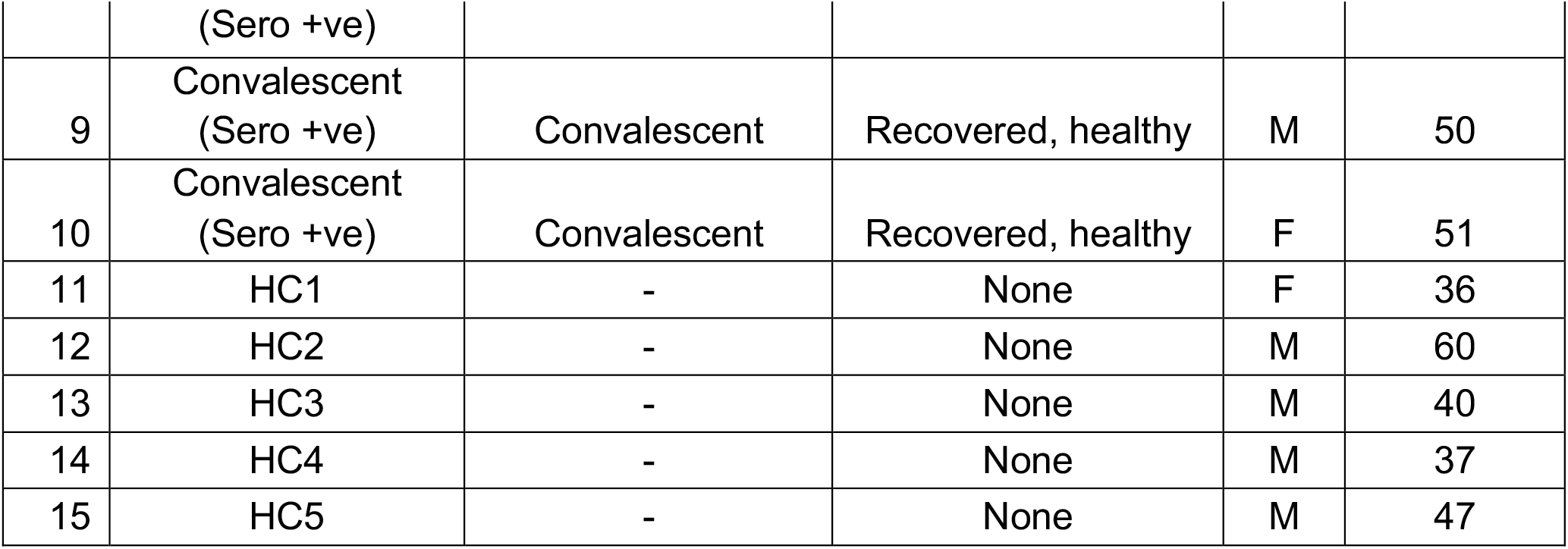
Patient demographics.

### Immunophenotyping of COVID-19 mild, moderate and convalescent COVID-19 patients

To compare the number of lymphocytes and monocytes amongst the four study groups, PBMCs were stained and analysed by flow cytometry. Based on live-cell percentage count (gating strategy; Suppl. Fig. 1a), both, lymphocytes and monocytes were not significantly different among mild, moderate and convalescent COVID-19 patients compared with HC (Suppl. Fig. 1b-d).

**Fig. 1:**
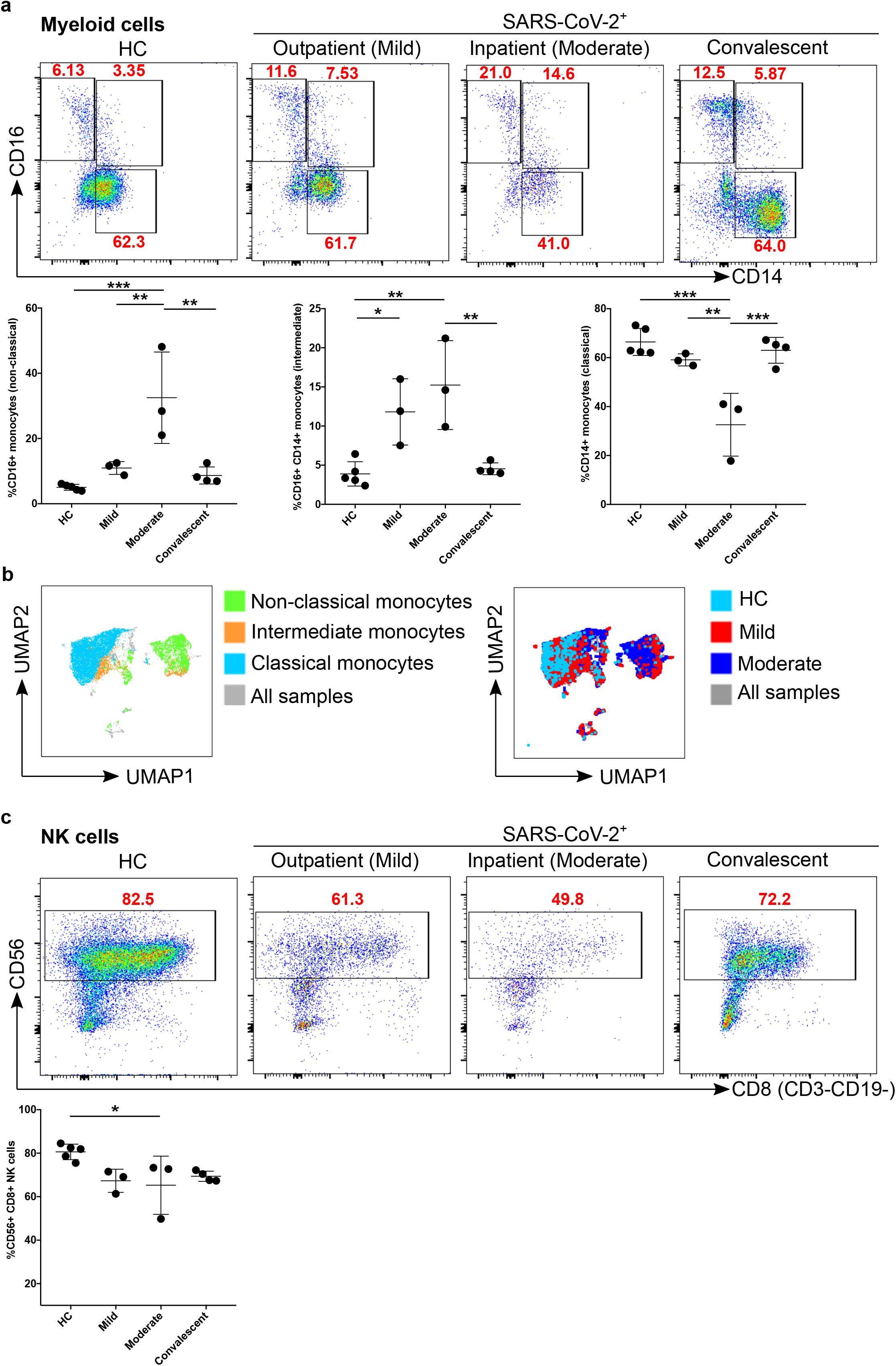
Comparison of monocytes and NK cell percentage amongst study groups. A. The stained PBMCs were gated on the monocyte population and CD3^+^CD19^+^ cells were excluded. Cell populations are displayed for CD16 and CD14 expression (upper FACS panel). One exemplary dot plot is shown per study group. The bar diagrams (lower panel) show the non-classical (CD16^+^CD14^-^), intermediate (CD16^+^CD14^+^) and classical (CD16^-^CD14^+^) monocytes. * P-value ≤ 0.05, * * P-value ≤ 0.01 and * * * P-value ≤ 0.001. B. UMAP plot for different subsets of monocytes (non-classical, intermediate and classical monocytes; left UMAP plot) from all the samples. Group comparisons among HC, mild and moderate (right UMAP plot). Colour coded information is provided for either different subsets of monocyte populations or patient groups. C. The stained PBMCs were gated on lymphocyte population and further excluded the CD3^+^CD19^+^ cells and examined for the CD56 and CD8 expression in HC, mild, moderate and convalescent (upper FACS panel). One exemplary dot plot is shown per study group. The bar diagram shows the CD56^+^CD8^+^CD3^-^CD19^-^ NK cells. * P-value ≤ 0.05.

### Increased inflammatory monocytes and reduced NK cells in moderate COVID-19 patients

Monocytes were further classified into classical, non-classical and intermediate based on the expression of CD16 and/or CD14 and were gated as described earlier^44^ (Suppl. Fig. 1a). We found that CD16^+^CD14^-^ patrolling (non-classical) monocytes were significantly increased (p=0.0004) in percentage in moderate patients compared to HC, whereas this number is decreased again significantly compared with convalescent patients (p=0.001) (Fig. 1a). The percentage of CD16^+^CD14^-^ (non-classical) monocytes was also significantly increased (p=0.006) in moderate patients compared with mild patients (Fig. 1a Panel I; left). Interestingly, CD16^+^CD14^+^ pro-inflammatory monocytes (intermediate) were again significantly increased in mild (p=0.03) and moderate (p=0.002) compared with HC as well as between moderate and convalescent (p=0.005) (Fig. 1a panel II; middle). Furthermore, we observed a significantly reduced percentage of CD14^+^CD16^-^ phagocytic monocytes (classical) in moderate compared with mild (p=0.004), HC (p<0.0002) and convalescent (p=0.0007) patients (Fig. 1a Panel III; right). Based on the results above using classical 2D flow cytometry analysis, we observed a major difference between HC *vs* mild or with moderate COVID-19 patient samples, therefore, we concatenated three groups together (equal number of cells from each sample; HC, mild and moderate) to identify the visual clustering of monocytes using Uniform Manifold Approximation and Projection (UMAP) for dimension reduction algorithm^45^ (Suppl. Fig. 2). Based on UMAP dimension reduction analysis (data-driven and gated cells), we observed a clear difference in the different subsets of monocytes between all three groups (Suppl. Fig. 2c, d & Fig. 1b). We observed that moderate COVID-19 patients mostly cluster in non-classical and partially in intermediate monocyte regions compared to HC, whilst mild COVID-19 patients cluster in all non-classical, intermediate and classical monocytes regions (Fig. 1b; right). We then explored the lymphoid cells compartment for NK cells (CD56^+^CD8^+^CD3^-^CD19^-^). We found that moderate patients were significantly different from HC (p=0.04). There was also a tendency of a decrease in NK cells in mild patients compared with HC (p=0.07), although, not reaching statistical significance (Fig. 1c).

**Fig. 2:**
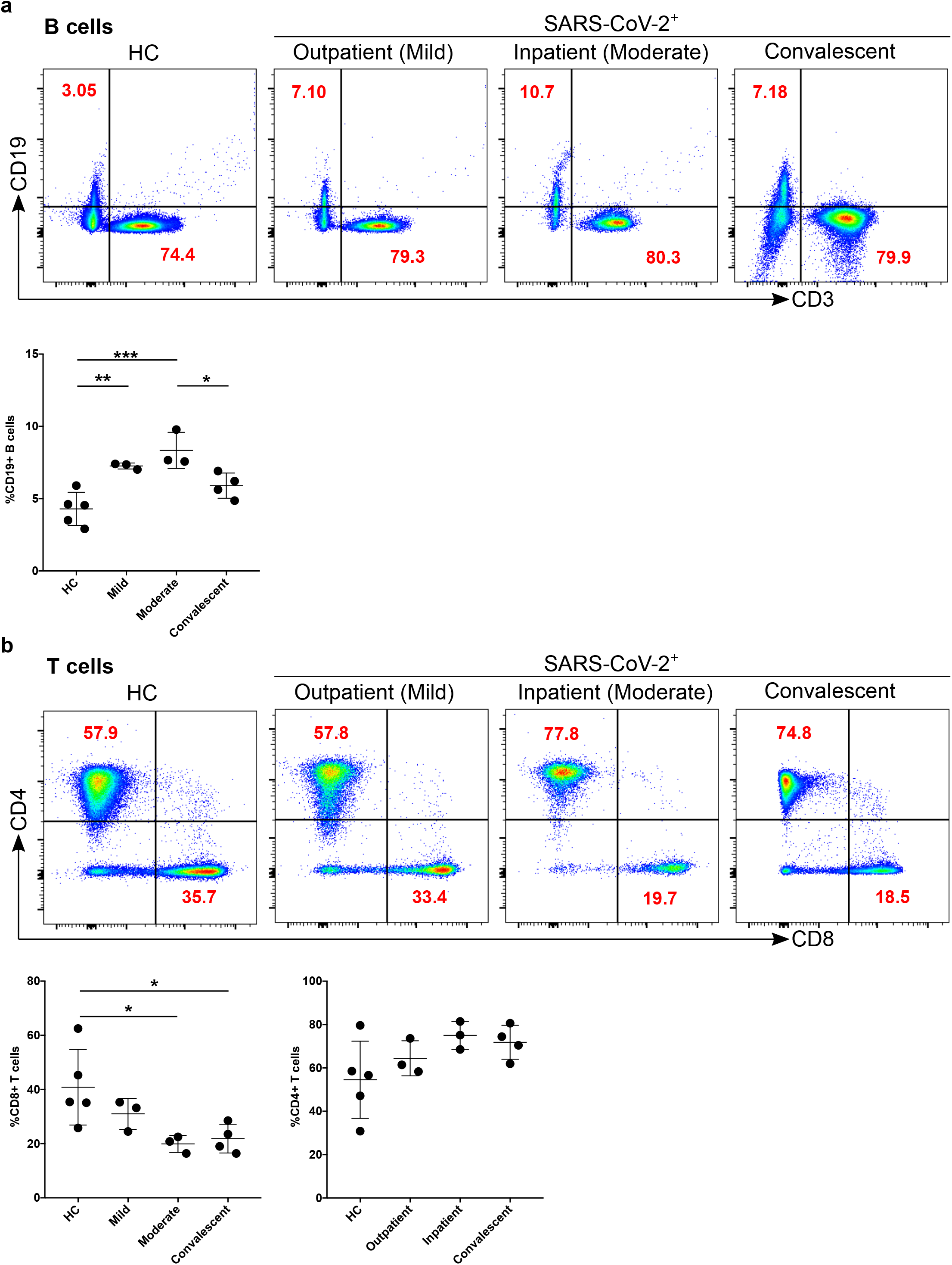
Increased B cells in mild and moderate patients and reduced CD8^+^ cytotoxic T cells in mild and convalescent patients A. The stained PBMCs were gated on lymphocyte population and examined for the CD19 and CD3 expression in HC, mild, moderate and convalescent (upper FACS panel). One exemplary dot plot is shown per study group. The bar diagram shows CD3^-^CD19^+^ B cells. * P-value ≤ 0.05, * * P-value ≤ 0.01 and * * * P-value ≤ 0.001. B. The CD19^-^CD3^+^ lymphocytes were examined for CD4^+^ and CD8^+^ T marker expression. One exemplary dot plot is shown per study group. There was statistically significant difference among HC, mild, moderate and convalescent (upper FACS panel). However, CD8^+^ T cells were significantly reduced in outpatient and convalescent patients. * P-value ≤ 0.05.

### Dynamics of B and T cells in mild, moderate and convalescent patients

Both T and B cells are indispensable for the immune response against viral infections including SARS-CoV-2. Firstly, we compared the number of B cells amongst the study groups, which give rise to virus-specific antibodies (see the gating strategy in Suppl. Fig. 1a). The CD19^+^CD3^-^ cells (B cells) were significantly increased in mild (p=0.008; fold change x1.7) and moderate (p=0.0008; fold change x1.9) patients compared with HC (Fig. 2a). Whilst, B cells were significantly decreased in moderate compared to convalescent (p=0.04) patients (Fig. 2a). Comparing CD3^+^CD19^-^ lymphocytes among the different patient groups we observed no significant difference. Moreover, CD3^+^ cells were analysed for the CD4^+^ and CD8^+^ T cell compartment. There was a tendency of increased CD4^+^ T cells for mild, moderate, and convalescent patients compared to HC, but no significant difference was observed among any of the groups. CD8^+^ T cells were significantly different between HC compared to moderate (p=0.04) or convalescent (p=0.04) patients (Fig. 2b). Finally, we characterized CD4^+^FOXP3^+^CD45R^-^ regulatory T cells (Tregs), however, no significant difference was observed among the different groups (Suppl. Fig. 3).

**Fig. 3:**
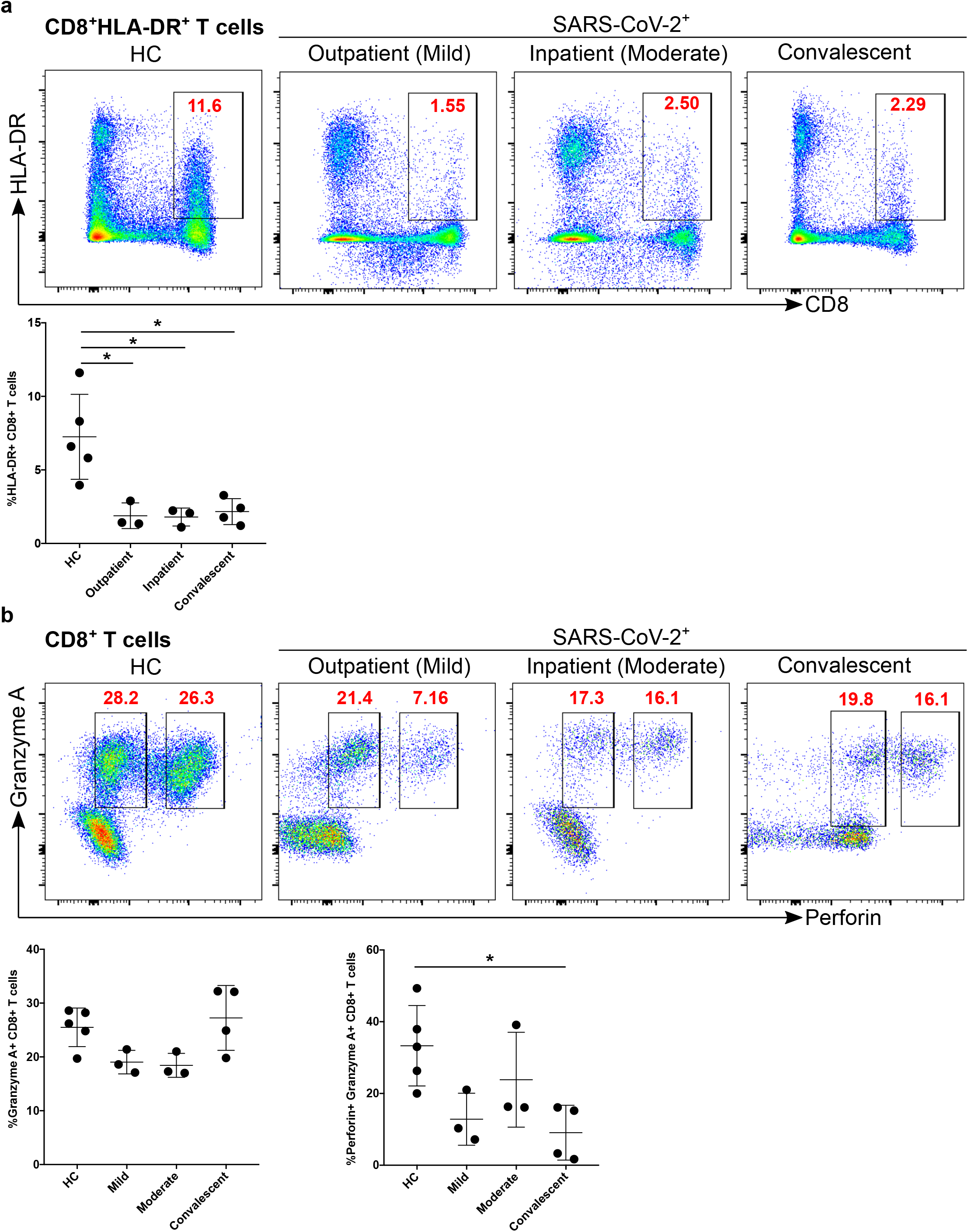
Decreased activation and cytotoxic functional protein expression of CD8^+^ T cells in convalescent patients A. CD8^+^ T cells were examined for the expression of activation marker HLA-DR (upper FACS panel). One exemplary dot plot is shown per study group. The bar diagram (lower panel) shows that HLA-DR was a significantly lower on CD8^+^ T cells in mild, moderate and convalescent COVID-19^+^ patients compared with HC. * P-value ≤ 0.05. B. CD8^+^ T cells were examined for the expression of their cytotoxic potential using granzyme A and perforin expression using IC staining (upper FACS panel). One exemplary dot plot is shown per study group. There was a statistically significant difference among HC, mild, moderate and convalescent (upper FACS panel) for granzyme A. The bar diagram (lower panel) shows that perforin expression was significantly lower on CD8^+^ T cells in convalescent COVID-19^+^ patients compared with HC, though mild and moderate represent a lower expression of perforin, but it did not to a significant level. * P-value ≤ 0.05.

### Impaired activation and defective cytotoxic functions of CD8^+^ T cells

We found that the percentage of CD8^+^ T cells were decreased in mild and convalescent patients compared to HC. Thus, we explored the activation status of CD8^+^ T cells based on HLA-DR expression. We found that CD8^+^ T cell activation status in all three groups of infected patients was significantly different from HC (mild p=0.01, moderate p=0.01, and convalescent p=0.01, Fig. 3a). We characterized the cytotoxic potential of CD8^+^ T cells based on granzyme A and perforin levels and found that there was a tendency of decreased granzyme A expression in mild, moderate and convalescent patients compared with HC (Fig. 3b), however, it did not reach significance. Perforin was significantly decreased in convalescent (p=0.02) patients compared with HC (Fig. 3b), although mild patients also had reduced levels (p=0.07), it did not reach statistical significance. Furthermore, we studied the expression of CD38, a marker of immune cell activation, which was significantly upregulated in convalescent patients compared with HC (p=0.04) (Fig. 4a). Similarly, convalescent patients had significantly increased numbers of PD-1^+^ CD38^+^ cytotoxic CD8^+^ T cells compared with HC (p=0.006), moderate (p=0.005) and mild (p=0.002), which reflects the exhaustion and non-responsiveness (anergy) of CD8^+^ T cells (Fig. 4b). Overall, our data suggested that CD8^+^ T cells have reduced activation, diminished expression of cytotoxic molecules such as perforin and granzyme A and have a severely exhausted phenotype.

**Fig. 4:**
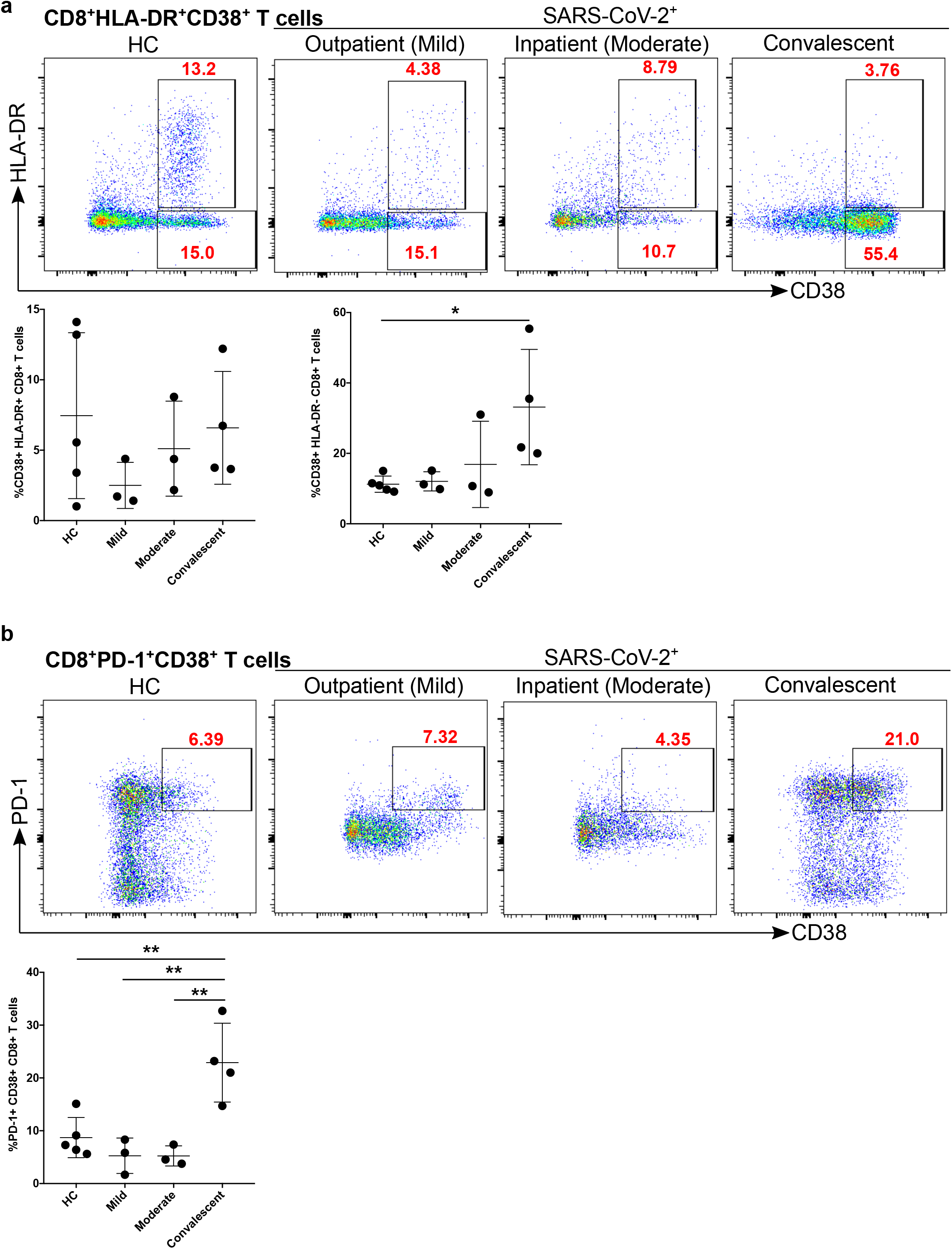
Increased exhausted CD8^+^ T cells in convalescent patients A. Expression of activation marker CD38 on CD8^+^ T cells (upper FACS panel). One exemplary dot plot is shown per study group. The bar diagram (lower panel) shows that CD38 expression was significantly higher on CD8^+^ T cells in convalescent COVID-19^+^ patients compared with HC. * P-value ≤ 0.05. B. Expression of activation marker CD38 and PD-1 on CD8^+^ T cells (upper FACS panel). One exemplary dot plot is shown per study group. The bar diagram (lower panel) shows that PD-1^+^CD38^+^ expression on was a significantly higher on CD8^+^ T cells in convalescent COVID-19^+^ patients compared with HC. * * P-value ≤ 0.01.

### Dynamics of metabolites production in the mild, moderate and convalescent patient

To establish a putative link between the metabolic state of immune cells and the impaired immune response, PBMCs from all patient groups were subjected to ^1^H-NMR spectroscopy analysis. We identified and quantified a total of 18 metabolites (Fig. 5a). Hereby, unsupervised Principal Component Analysis (PCA) showed that spectral data from mild and moderate patients formed overlapping clusters. However, HC and convalescent patients clustered together (Fig. 5b), indicating a strong difference in metabolite levels between an infectious state compared to healthy or recovered groups. Statistical analysis of the four different groups revealed that 15 metabolites showed p-values < 0.05, with the highest significance for metabolites related to energy metabolism (Fig. 5c, Suppl. Fig. 3 & Table 2). The data indicate that during infection, there is a strong consumption (reflecting reduced levels) of glucose, acetate and formate, whilst lactate levels are increased. Furthermore, we also found very high levels of fructose in PBMCs from mild patients, medium concentrations in moderate and, low levels in HC and convalescent patients (Fig. 5c). Furthermore, glutamate was almost abolished in mild and moderate patients, potentially as a consequence of enhanced production of α-ketoglutarate in the TCA cycle in PBMCs *via* glutamate dehydrogenase (Fig. 5c). Levels of other amino acids such as glycine and isoleucine were low in mild and moderate patients compared with HC, while creatine and alanine were high in mild patient and moderate patient respectively (Fig. 5c).

**Table 2:** Summary of metabolites dysregulated in PBMCs.

| No | Metabolites | HC | Mild | Moderate | Convalescent |
| --- | --- | --- | --- | --- | --- |
| 1 | Glucose | ↑↑ | ↓↓ | ↓↓ | ↑↑ |
| 2 | Formate | ↑↑ | ↓↓↓ | ↓↓ | ↑↑ |
| 3 | Acetate | ↑ | ↓↓ | ↓ | ↑ |
| 4 | Lactate | ↓↓ | ↑↑ | ↑↑ | ↓↓ |
| 5 | Fructose | ↓ | ↑↑ | ↑ | ↓ |
| 6 | Glutamate | ↑ | ↓ | ↓ | ↑ |
| 7 | Citrate | ↓ | ↑↑ | ↑ | ↓ |
| 8 | Taurine | ↓↓ | - | ↑↑ | ↓ |
| 9 | Creatine | ↓ | ↑ | - | - |
| 10 | Alanine | ↓ | - | ↑↑ | ↓ |
| 11 | Glycine | ↑ | ↓ | ↓ | - |
| 12 | Isoleucine | - | ↓↓ | ↓ | ↑ |

**Fig. 5:**
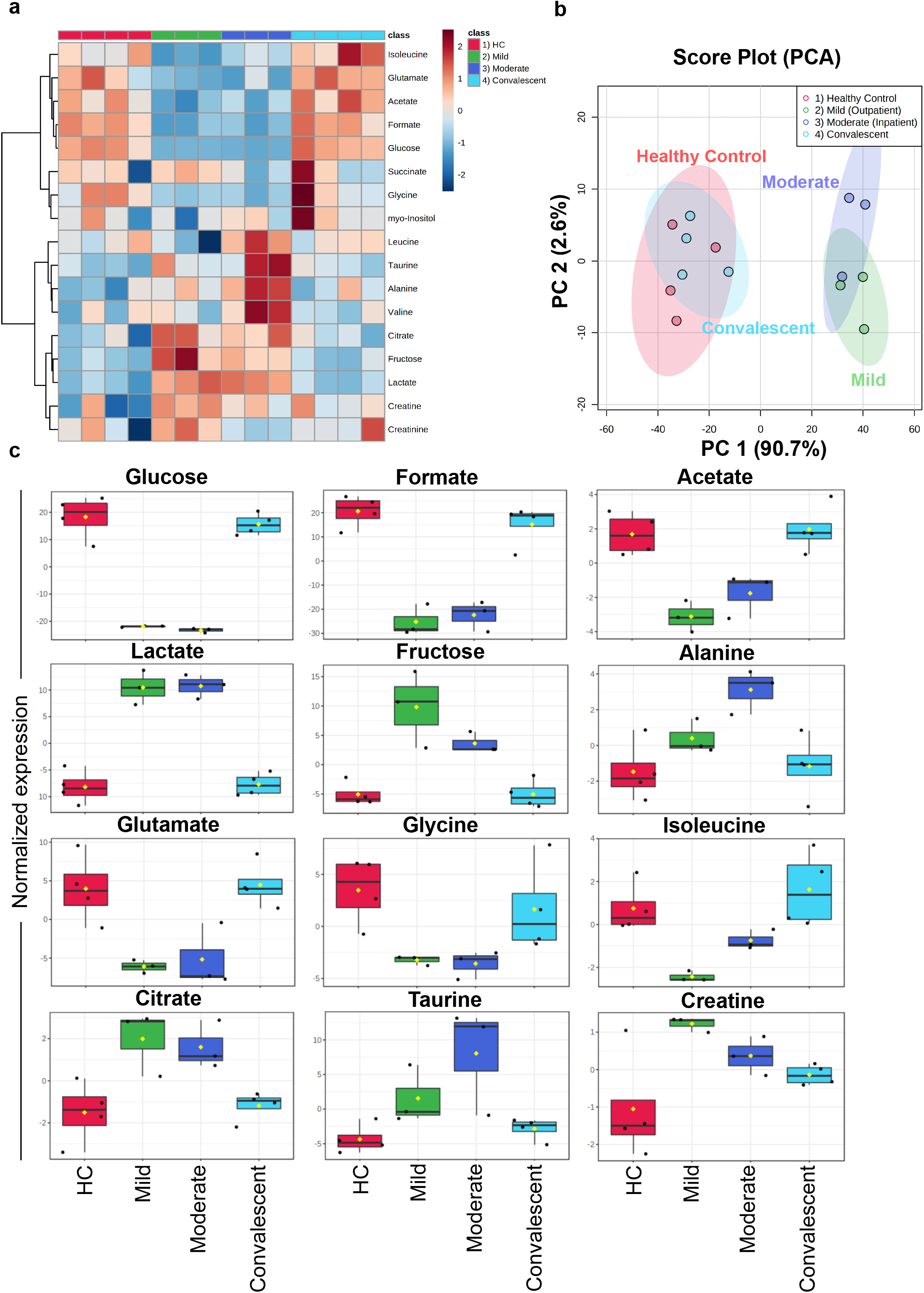
^1^H-NMR spectroscopy of PBMC extracts A. Heatmap of featured metabolites’ concentrations plotted with SARS-CoV-2 progression group clustering. B. Principle component analysis (PCA) was performed to identify the clustering of two different groups. HC and convalescent COVID-19 patient samples cluster together while SARS-Co-2 infected mild and moderate patients cluster in a separate cluster with PC1: 90.7% and PC2: 2.6%. C. Box plots for differentially abundantly present metabolites in the different group including HC, mild, moderate, and convalescent COVID-19 patient. * P-value ≤ 0.05, * * P-value ≤ 0.01 and * * * P-value ≤ 0.001.

To find an association between different metabolites, we applied the variable importance of projection (VIP) score. We found that formate and glucose had the highest score compared to other metabolites (Fig. 6A). To determine if additional metabolites are positively associated with changes in glucose, lactate and fructose, we performed a pattern hunter analysis for all metabolites. We found that high glucose levels correlated with high formate, acetate and glutamate and low lactate and fructose (Fig. 6b), indicating enhanced glycolysis and TCA cycle in PBMCs. Similarly, fructose, which is entered *via* fructose-1-phosphate and dihydroxyacetone phosphate (DAP) into the glycolysis, is correlated positively with lactate and citrate and a decrease in acetate and formate, respectively (Fig. 6b). Interestingly, levels of the ROS scavenger taurine are only positively correlated with lactate and fructose, but not glucose (Fig. 6).

**Fig. 6:**
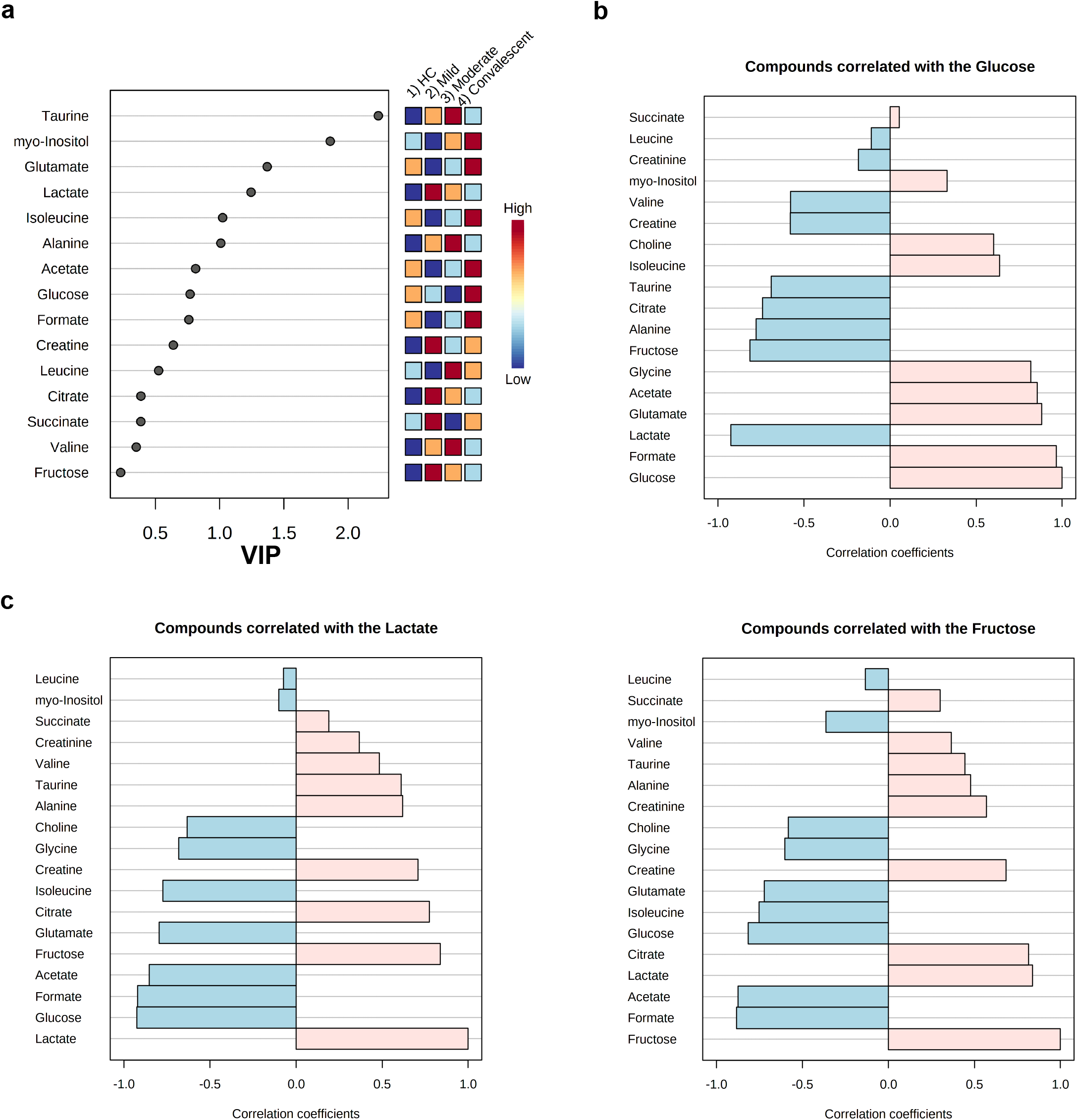
Pattern hunter plots provide an insight of close correlations with other metabolites during COVID-19 infection. A. Variable Importance in Projection (VIP) scores for all metabolites in the four studied groups. B. Pattern hunter plot for glucose. C. Pattern hunter plot for lactate and fructose.

## Discussion

SARS-CoV-2 infections are an intense and rapidly evolving area of research due to the ongoing global pandemic^14,25^. In this study, we used flow cytometry and ^1^H-NMR to decipher the cell proportions and functional state of immune cells (PBMCs) in mild, moderate and convalescent COVID-19 patients compared to HC. Recent reports from COVID-19 patients suggested that mild and severe patient had lymphopenia^11,46-48^. Here, we found that the myeloid cell compartment in PBMCs based on CD16 and CD14 markers suggested that the percentage of non-classical and intermediate monocytes were increased during an active mild or moderate SARS-CoV-2 infection, once infections are cleared the monocyte percentage numbers return to normal. These results are in accordance with some of the recently published studies^49-51^.

In our cohort, specifically CD56^+^CD8^+^NK cells were decreased during active SARS-CoV-2 viral infections (moderate), while during recovery the numbers were comparable to HC as reported by others^52^. Similarly, another recent study supports our finding by showing the decrease in the number of NK cell subsets in COVID-19 patients, with no change in CD56^bright^ or CD56^dim^ cells^53^. Furthermore, based on single-cell RNA-sequencing data, a reduced number of NK cells was reported in the recovered COVID-19 patients^48^. Thus, altogether, it appears that CD56^+^NK cells and their subsets could have an important function during the ongoing and clearance of SARS-CoV-2 infections. However, further validation studies are warranted using different *in vivo* model systems with appropriate control groups to pinpoint the exact role of different NK cell subsets in SARS-CoV-2 infection. CD19^+^ B lymphocytes were increased during infection and remain slightly higher than HC, thus reflecting the antibody response against the SARS-CoV-2 virus. Thus, our data suggest that these patients were able to generate SARS-CoV-2 specific B cells, this needs further scientific validation.

A major difference was found in the T lymphocytes compartment. CD8^+^ T cells were significantly decreased in moderate and convalescent patients as reported earlier^52^. Thus, it appears that during viral infection non-virus specific CD8^+^ T cells undergo apoptosis, whilst the viral-specific surviving CD8^+^ T cells are clonally expanded but appeared to have lost their effector functions^54^. To confirm this, we first measured the activation status of CD8^+^ T cells and found that CD8^+^ T cells appeared to be less activated based on their HLA-DR activation marker^26^. Further, CD8^+^ T cells were examined for additional activation marker CD38, which is involved in cell adhesion, signal transduction and calcium signalling^55^, was found to be upregulated in convalescent patients but not during active infection. These CD38^+^CD8^+^ T cells also expressed higher levels of PD-1, which is an immune checkpoint and marker of exhaustion^24,30,56-58^. PD-1 further guards against autoimmunity promote apoptosis of antigen-specific T cells and promotes self-tolerance by suppressing T cell inflammatory activity^59^. Thus, viral infection leaves convalescent patients with exhausted phenotypes. We uncovered that although there was not a significant change in the numbers of Tregs in COVID-19 patients, there was a trend towards elevated levels of Tregs in COVID-19 patients and rescued Tregs in convalescent patients, in agreement with the previous studies^56^.

A key finding of our study was the surprising observation that granzyme A and perforin secreting CD8^+^ T cells were significantly reduced in convalescent patients. These results are in agreement with the recent single-cell transcriptomics analysis which suggests that granzyme B and perforin-1 transcripts were also decreased in convalescent COVID-19 patients compared with moderate or severe illness^60^. In contrast to our results, in young patients, granzyme A or B and perforin levels were increased in mild and moderate cases. Conversely, in elderly COVID-19 patients, there was a reduced expression of granzyme A and perforin^61^. Another study suggested that decreased perforin and granzyme A levels in CD4^+^ T cells, CD8^+^ T cell and NK cells is associated with severely afflicted COVID-19 patients^62^. Furthermore, single-cell transcriptomics analysis of SARS-CoV-2 reactive CD8^+^ T cells in exhausted and non-exhausted subsets were analyzed^63^. These exhausted CD8^+^ T cells were increased in frequency and displayed lesser cytotoxicity and inflammation features in mild COVID-19 patients compared to severe patients^63^. A genetic study performed on two cases highlights the importance of the perforin gene variant A91V which results in the rapid demise in young COVID-19 patients^64^. The possible implication of our finding is that convalescent patients, specifically including cancer patients under treatment, could be susceptible to future opportunistic infections with other viruses including different variants of SARS-CoV-2. However, further, a large cohort study is warranted to understand the potential functions of these molecules (granzyme or perforin) in protection or susceptibility against the COVID-19 infection at gene and function levels.

To date, the general metabolic physiology of PBMCs is not well defined in the literature. However, it is clear now that PBMCs are dependent on circulating nutrients and hormones in the blood^65^. A defective immune response in COVID-19 patients prompted us to investigate the metabolic functions of these immune cells. Our metabolomics data indeed shows that PBMCs from actively infected patients have a distinct metabolic profile from convalescent or healthy individuals. The most notable difference we observed was for metabolites from the glycolysis and oxidative phosphorylation (TCA cycle) pathway, which is per recently published transcriptome data for PBMCs^39,43^. Metabolites such as glucose, formate, acetate and choline were also reduced in PBMCs in infected patients whereas, HC and convalescent patients had a normal profile. Accordingly, the glycolytic pathway end products such as lactate were higher in active mild and moderate COVID-19 patients compared with HC and convalescent individuals. Therefore, our data suggest that PBMCs (which constitute a major fraction of T lymphoid cells: 70-80%) may have modulated their metabolic functions, particularly favouring the oxidative phosphorylation pathway over the glycolytic pathway, to meet the high demands of energy needed to combat the ongoing viral infection.

A recent report suggested that elevated glucose levels enhance SARS-CoV-2 replication and cytokine expression in monocytes and glycolysis sustains the viral-induced monocyte response^66^. Recently, it was emphasized that glucose consumption in PBMCs during COVID-19 disease could be also a read-out of cytokine storms^34^. Further, a higher abundance of citrate in PBMCs suggested that perhaps T cells could use the oxidative phosphorylation pathway for energy consumption to endure the infection, as recent transcriptomic data also suggested that higher expression of genes related to oxidative phosphorylation both in peripheral mononuclear leukocytes and bronchoalveolar lavage fluid (BALF) could play a crucial role in increased mitochondrial activity during SARS-CoV-2 infection^34^.

Another remarkable finding of our study was the increase of fructose levels in PBMCs during the course of infection. Previous findings suggested that fructose is involved in the inflammatory pathways for the production of IL-1β and IL-6 production^67^. Thus, it is conceivable that the immune cells (most probably monocytes) could be triggered by higher fructose and simultaneously induce inflammation and IFN-γ production by T cells^67^. These findings correlate with recent transcriptomic studies on the BALF from infected COVID-19 patients and plasma of COVID-19 patients that also identified changes in fructose metabolism^34,68^.

Several amino acids are involved in anti-inflammatory effects, especially arginine, glutamine and glycine appeared to improve lung damage induced by infections^69^. Administration of glutamine reduced the inflammatory cytokines, whereas arginine or glycine reduced IL-6 and CXCL-1 expression in the alveolar epithelium^70^. Additionally, glutamine is an important signalling molecule involved in activating mammalian target of rapamycin (mTOR) signalling which is critical for immune cell activity and inhibiting catabolic functions such as protein degradation and apoptosis^71^. In our study, we found a reduced abundance of glutamate and glycine in PBMCs during the mild and moderate COVID-19 patients. Reduced glucose and glutamine are involved in the hexosamine biosynthetic pathway and could be responsible for poor T cell clonal expansion and function^41^. The exact molecular mechanism of individual amino acids is a complex phenomenon and currently unclear. However, at least in theory, supplementation of glutamine and glycine could be beneficial for COVID-19 patients.

We finally observed a reduction of granzyme A and perforin levels in CD8^+^ T cells and detected the ambient level of antioxidant amino acid taurine in the convalescent patients, which could be involved in the modulation of cytotoxic functions of CD8^+^ T cells. Both granzyme A and perforin are involved in ROS production and taurine serves as ROS scavenger^72,73^. Thus, decreased granzyme A and perforin could be implicated in reduced ROS production for the impaired effectiveness of CD8^+^ T cells in either convalescent patients or COVID-19 patients. This should be the case, as taurine levels are generally increased during active infection in mild/moderate patients compared to healthy controls and are not specifically decreasing due to granzyme A and perforin lacking ROS activity in COVID-19 patients. However, this finding needs further investigation to validate this hypothesis as it is unclear how ROS and taurine act together to affect the cytotoxic functions of immune CD8^+^ T cells. In summary, the metabolomics data generated in this study provides first and crucial insights into the complex metabolic changes of PBMCs during SARS-CoV-2 infections, warranting further future in-depth investigation.

## Conclusions

Using immunophenotyping and metabolomics approaches we detected significant changes in PBMC samples of mildly and moderately affected COVID-19 as well as convalescent patients compared with healthy controls. The reduced percentage of NK cells in both mild and moderate patient groups corresponded with the clustering of PBMCs metabolite levels in the principal component analysis distinct from the cluster formed by healthy and convalescent individuals. The dramatically changed metabolic activity and pathways, such as glycolysis and TCA cycle, might not only lead to a vulnerability of COVID-19 patients to subsequent infections but can also offer insights into how PBMCs could be manipulated towards a better survival and personalized treatment of moderate and severe COVID-19 patients.

## Limitation of the study

The current study is performed from the samples obtained during the first wave of infections and suffers from the limited number of patients used in the study. The use of appropriate controls such as influenza viral infection would have been useful for a more generalized conclusion. Therefore, results obtained in future studies might differ from subsequent waves of infections in new patient cohorts. Further, the novel SARS-CoV-2 variants or vaccine candidates might promote alternative host immune evasion strategies to infect the host, thus, eliciting a different immune response. Nonetheless, this study certainly highlights the importance of distinct cell types as well as the crucial function of cell metabolism during SARS-CoV-2 viral infection and even after recovery.

## Materials and Methods

### Ethics statement

The study protocols were approved by the University of Tübingen, Germany Human Research Ethics Committee (TÜCOV: 256/2020BO2 (convalescent study), COMIHY: (225/2020AMG1) (outpatient study)-COMIHY, EUDRA-CT: 2020-001512-26, ClinicalTrials.gov ID: NCT04340544, and COV-HCQ: (190//2020AMG1) (inpatient study)-COV-HCQ, EUDRA-CT: 2020-001224-33, ClinicalTrials.gov ID: NCT04342221, 556/2018BO2) and all associated procedures were carried out in accordance with approval guidelines. All participants provided written informed consent in accordance with the Declaration of Helsinki.

### Study participants

SARS-CoV-2 positive patients were used for this study and no other virus species were analysed in this study (COMIHY and COV-HCQ). Blood was collected from COVID-19 patients enrolled into two different prospective randomized, placebo-controlled, double-blind clinical trials evaluating the safety and efficacy of hydroxychloroquine in COVID-19 outpatients (COMIHIY) and hospitalized patients (COV-HCQ). We analysed subsets of these study cohort and used outpatient (n=3; COMIHY) which came to a specified outpatient ward at the Institute of Tropical Medicine with mild symptoms and blood was taken and usually defined as D1 outpatients. Inpatients (n=3; COV-HCQ), blood was taken after 7-9 days after study inclusion defined as D7. These patients had moderate symptoms needing hospital care, however not being transferred to the intensive care unit in the hospital. Furthermore, convalescent COVID-19 patients (n=4) were defined as positive for serum antibody reactive to SARS-CoV-2 and blood was taken when they visited the Institute of Tropical Medicine for testing of antibody levels. Amongst this cohort, 3 persons reported mild fever for 10-11 days and 1 individual reported no fever but found positive for SARS-CoV-2 antibodies. Blood from healthy controls (n=5) was obtained from the hospital blood bank.

### Flow cytometry and UMAP data analysis

PBMCs were isolated by the standard Ficoll method^74^. A total of 1-2 ×10^6^ PBMCs per participants were used for three FACS panels (Table 2). In brief, firstly, to distinguish between live from dead, the cells were incubated with LIVE/DEAD Fixable Infra-Red Dead stain (Thermofisher) for 15 minutes at room temperature (RT) into 1:40 diluted dye in DPBS. Subsequently, after LIVE/DEAD staining, cells were stained with surface markers in DPBS (Sigma) with Super Bright stain Buffer (Thermofisher) for 30 minutes at RT. After surface staining cells were also stained for intracellular (IC) markers. Before IC staining, cells were fixed for 30-45 minutes and permeabilized for 5 minutes followed by IC antibody incubation for additional 30 minutes at RT. Cells were washed and resuspended in DPBS containing 2%FBS. Fixing of cells was performed irrespective of whether the panel was used for IC staining or not to prevent the possible contamination during the acquisition of the samples. All the staining procedures were performed in the dark to avoid the photo-bleaching of dyes. For each sample, 200,000 cells were acquired using BD LSRFortessa (core facility) equipped with 4 lasers (violet, blue and yellow-green and Red). Data were analysed using Flow Jo (Tree Star) and fluorescence minus one controls (FMO) were used for setting up the arbitrary gates for the major cell markers. Furthermore, UMAP dimensional reduction analysis was performed using UMAP plug-in^45^ in Flowjo software using default setting except for minimum distance (0.2 instead of 0.5) and population (n=15) as advised by a plug-in. First, dead/debris was removed by gating (FSC-A vs SSC-A; linear scale) then using FSC-A vs FSC-W, we focussed on singlets. The singlets were again gated for live and dead discrimination. An equal number of live cells from each sample (HC, mild and moderate) were concatenated and exported as a single FCS file. This FSC file was subjected to UMAP analysis, each cell population was either monocytes or lymphocytes were again subject to subsequent UMAP analysis for clustering of specific sub-populations. Based on gating or antibodies-stained cells (data driven) analysis was performed and summarized in Suppl. Fig. 2.

**Table 2:** Antibodies and other reagents used for Flow cytometry.

| No. | Product Name | Clone | Fluorochrome | Product ID | Company |
| --- | --- | --- | --- | --- | --- |
|  |  | **NK cells and Monocytes (Panel 1)** |  |  |  |
| 1 | CD3 | UCHT1 | eFluor 450 | 48-0038-42 | Thermofisher |
| 2 | CD4 | SK3 | SuperBright 600 | 63-0047-42 | Thermofisher |
| 3 | CD8a | SK1 | PerCP-eFluor 710 | 46-0087-42 | Thermofisher |
| 4 | CD19 | HIB19 | eFluor 506 | 69-0199-42 | Thermofisher |
| 5 | CD45-RA | HI100 | PE-Cy7 | 25-0458-42 | Thermofisher |
| 6 | HLA-DR | L243 | Alexa Fluor647 | A51010 | Thermofisher |
| 7 | CD38 | HIT2 | PE-eFluor610 | 61-0389-42 | Thermofisher |
| 8 | CD56 | MEM188 | PE | MA119638 | Thermofisher |
| 9 | CD16 | 3G8 | Super Bright702 | 67-0166-42 | Thermofisher |
| 10 | CD14 | 61D3 | Alexa Fluor700 | 56-0149-42 | Thermofisher |
| 11 | Foxp3 (IC) | PCH101 | FITC | 11-4776-42 | Thermofisher |
|  |  | **CD8 exhaustion, T helper follicular cells (Tfh) and antibody-secreting cell (ASC) (Panel 2)** |  |  |  |
| 1 | CD3 | UCHT1 | eFluor 450 | 48-0038-42 | Thermofisher |
| 2 | CD19 | HIB19 | eFluor 506 | 69-0199-42 | Thermofisher |
| 3 | CD4 | SK3 | Super Bright 600 | 63-0047-42 | Thermofisher |
| 4 | CD8a | SK1 | PerCP-eFluor 710 | 46-0087-42 | Thermofisher |
| 5 | CD38 | HIT2 | PE-eFluor 610 | 61-0389-42 | Thermofisher |
| 6 | CD27 | O323 | Alexa Fluor700 | 56-0279-42 | Thermofisher |
| 7 | CXCR5 (CD185) | MU5UBE E | FITC | 11-9185-42 | Thermofisher |
| 8 | ICOS (CD278) | C398.4A | PE | 12-9949-81 | Thermofisher |
| 9 | PD-1 (CD279) | eBioJ105 (J105) | PE-Cy7 | 25-2799-42 | Thermofisher |
| 10 | HLA-DR | L243 | Alexa Fluor647 | A51010 | Thermofisher |
|  |  | **Cytotoxic potential** |  |  |  |
| 1 | CD4 | SK3 | SuperBright600 | 63-0047-42 | Thermofisher |
| 2 | CD8 | SK1 | PerCP-eFluor710 | 46-0087-42 | Thermofisher |
| 3 | CD19 | HIB19 | eFluor 506 | 69-0199-42 | Thermofisher |
| 4 | CD38 | HIT2 | PE-eFluor 610 | 61-0389-42 | Thermofisher |
| 5 | HLA-DR | L243 | Alexa Fluor 647 | A51010 | Thermofisher |
| 6 | GZMA (IC) | CB9 | PE | 12-9177-42 | Thermofisher |
| 7 | GZMB (IC) | GB11 | Alexa Fluor 488 | MA5-23639 | Thermofisher |
| 8 | Perforin (IC) | dG9 | PE-Cy7 | 12-9177-42 | Thermofisher |
|  |  | **Other Flow reagents** |  |  |  |
| 1 | Ultracompensation bead |  |  | 01-2222-42 | Thermofisher |
| 2 | Live/Dead Fixable Near IR staining kit |  |  | L10119 | Thermofisher |
| 2 | FOXP3/TRN FACTOR STAIN BUFFER SET |  |  | 00-5523-00 | Thermofisher |
| 3 | FLOW STAIN BUFFER SOLN |  |  | 00-4222-57 | Thermofisher |
| 4 | SB COMPLETE STAINING BUFFER |  |  | SB-4401-42 | Thermofisher |
| 5 | DPBS |  |  | D8537 | Sigma |
| 6 | Pancoll human |  |  | P04-601000 | Pan Biotech |

### ^1^H-NMR metabolomics

To obtain PBMCs metabolites, PBMCs were suspended in an optimized solvent extraction mixture of 9:1 (methanol: chloroform) as described elsewhere in detail^75^ and extracted with a focused ultrasound system (Covaris E220, Woburn, USA). The extraction solutions were evaporated to dryness for 4 hours in a vacuum concentrator and afterwards pellets resuspended with 45 µL in a 1 mM TSP containing deuterated phosphate buffer. After centrifugation at 20,000 x g for 10 min to remove residual macromolecules, 40 µL of the clear supernatant were transferred to 1.7 mm NMR tubes. Spectra were recorded on ultrashielded 600 MHz spectrometer (Bruker AVANCE III HD, Karlsruhe, Germany) with a triple resonance 1.7 mm room temperature probe. Spectra used for analysis were acquired with a 2h 55min lasting CPMG pulse program. Metabolite annotation and quantification were done with ChenomX NMR Suite 8.3.

### Statistical analysis

Bar diagrams were prepared using GraphPad Prism 6.0. Data shown are means ± SD. FACS data were analysed using one-way analysis of variance (ANOVA) for multiple group comparisons (mild, moderate, convalescent and HC) in GraphPad Prism software. No matching or pairing was used. Assumed Gaussian distribution with equal standard deviations (SDs) for experimental design. The mean of each group was compared with the mean of every other group and Tukey’s post-hoc tests for multiple comparisons were employed. P-value considered significantly less than 0.05 or equal. Metabolite concentrations from ^1^H-NMR analysis were exported as comma-separated value spreadsheet file to MetaboAnalyst 4.0 software, normalized with probabilistic quantile normalization (PQN) and range scaled. Unsupervised principal component analysis (PCA), a dimensionality-reduction method was used for clustering of all metabolites (PC1 and PC2 only). Multiple groups comparison was performed using one way ANOVA. Multivariate data analysis techniques such as partial least squares analysis (PLSDA) and variable’s importance in the PLSDA (VIP) score analysis were used to find the correlation among different metabolites and various groups. Hierarchical clustering of all the metabolites was measured and distance measurement was performed with Pearson r correlation coefficient.

## Figure legends

**Suppl. Fig. 1:** Total % counts of monocytes and lymphocytes from PBMCs of COVID-19 patients.

A. Fixed PBMCs samples (1×10^6^ cells) were acquired on flow cytometry on 2-3 different days for the entire experiments. Total 200,000 cells were acquired by flow cytometry and gating was performed based on FSC and SSC parameters for lymphocytes, monocytes and dead cells as described earlier^76-78^. Gating strategy for T lymphocytes (CD3, CD4 and CD8) monocytes (CD14 and CD16)^44^, NK cells (CD56) using FMO controls.

B. The bar graphs represent the % of lymphocytes and monocytes.

**Suppl. Fig. 2:** UMAP analysis of lymphocytes and monocytes cell populations.

A. UMAP pseudo-colour plot was derived from all the samples. All the FCS files (used only live-cell gated population; minimum cells per samples were 25,000) were concatenated from HC, mild and moderate patients (upper panel). Representation of lymphocytes and monocytes populations in UMAP plot (middle panel). Presentation of different lymphocytes subpopulations like CD4^+^ T cells, CD8^+^ T cells, B cells, NK cells, NKT cells and regulatory T cells (lower panel).

B. UMAP plot for overlaying of different groups.

C. UMAP plot for different subsets of monocytes.

D. UMAP plots for individual antibody (protein) expression on monocyte populations. Scale bar shows the expression level of protein expression on the cell clusters (blue -lowest and red highest expression; relative expression).

**Suppl. Fig. 3:** Kinetics of regulatory T cells is not affected significantly in mild, moderate and convalescent patients.

Foxp3^+^ expression on CD19^-^CD3^+^CD4^+^CD45RA^-^ T cells to identify the regulatory T cells in HC, outpatient, outpatient and convalescent (upper FACS panel). There was a statistically significant difference among HC, mild, moderate and convalescent (upper FACS panel).

**Suppl. Fig. 4:** Metabolite analysis in COVID-19 patients.

A. Analysis of Variance (ANOVA) for multi-group comparisons.

B. Partial Least Squares Discriminant Analysis (PLS-DA) scores plot.

C. Hierarchical clustering of metabolites (distance measured with Pearson r correlation coefficient).

D. Boxplots for branched-chain amino acids valine and leucine.

## Author’s contributions

YS: Overall study design and project coordination, flow cytometry experiments, data analysis and interpretation and initial metabolic sample preparation, funding generation, and writing the manuscript.

CT: Metabolites sample preparation, Processing of the sample on ^1^H-NMR, data analysis, data interpretation and writing

MO: Metabolites sample preparation, Processing of the sample on ^1^H-NMR

RF, NK: Provided patient materials, Isolation of PBMCs, Performed the experiments for flowcytometry in BSL-2 facility, flow data interpretations.

RB: Isolation of PBMCs from HC and sample preparation for flow cytometry. SO, MS, NC, OR: Study design, providing research tools, funding generation, data interpretation and writing and amending the manuscript.

All the authors have seen the manuscript, substantially contributed and agreed to be co-author.

## Acknowledgements

We thank you all the patients who participated in this study and FACS core facility (Klinikum-Berg) for accessing the Flow cytometry for the experiments. We also thanks Jana Held and Andrea Kreidenweiss from Institute of Tropical Medicine, Tübingen university for their help with coordination with clinical Trials and proof-reading of the manuscript. We acknowledge support by Deutsche Forschungsgemeinschaft (DFG) and Open Access Publishing Fund of University of Tübingen.

## Funding

RB is supported by Deutsche Forschungsgemeinschaft (DFG Project no. 426724658). EKFS (MS), DFG (OR), Ferring Pharmaceutical (YS and MSS). To MSS the Margarete von Wrangell (MvW 31-7635.41/118/3) habilitation scholarship co-funded by the Ministry of Science, Research and the arts (MWK) of the state of Baden-Württemberg and by the European Social Funds. Funders have no role in the study design and data analysis. To RF the German Federal Ministry of Education and Research (BMBF) (BMBF-01KI2052) and the German Federal Ministry of Health (BMG) (BMG-ZMVI1-1520COR801).

## Competing interests

Authors declare no financial competing interests.

### DECOI Consortia

The members of the Deutsche COVID-19 Omics Initiative (DeCOI) are Janine Altmüller, Angel Angelov, Robert Bals, Alexander Bartholomäus, Anke Becker, Daniela Bezdan, Ezio Bonifacio, Peer Bork, Nicolas Casadei, Thomas Clavel, Maria Colome-Tatche, Andreas Diefenbach, Alexander Dilthey, Nicole Fischer, Konrad Förstner, Sören Franzenburg, Julia-Stefanie Frick, Gisela Gabernet, Julien Gagneur, Tina Ganzenmüller, Marie Gauder, Alexander Goesmann, Siri Göpel, Adam Grundhoff, Torsten Hain, Andrè Heimbach, Michael Hummel, Thomas Iftner, Angelika Iftner, Stefan Janssen, Jörn Kalinowski, René Kallies, Birte Kehr, Andreas Keller, Sarah Kim-Hellmuth, Christoph Klein, Oliver Kohlbacher, Karl Köhrer, Michael Knop, Jan O. Korbel, Peter G Kremsner, Denise Kühnert, Ingo Kurth, Markus Landthaler, Yang Li, Kerstin Ludwig, Oliwia Makarewicz, Manja Marz, Alice McHardy, Christian Mertes, Sven Nahsen, Markus Nöthen, Francine Ntoumi, Peter Nürnberg, Uwe Ohler, Stephan Ossowski, Jörg Overmann, Silke Peter, Klaus Pfeffer, Anna R. Poetsch, Alfred Pühler, Nikolaus Rajewsky, Markus Ralser, Olaf Rieß, Stephan Ripke, Ulisses Nunes da Rocha, Philip Rosenstiel, Antoine-Emmanuel Saliba, Leif Erik Sander, Birgit Sawitzki, Philipp Schiffer, Wulf Scheider, Eva-Christina Schulte, Joachim L. Schultze, Alexander Sczyrba, Yogesh Singh, Michael Sonnabend, Oliver Stegle, Jens Stoye, Fabian Theis, Janne Vehreschild, Thirumalaisamy P Velavan, Jörg Vogel, Max von Kleist, Andreas Walker, Jörn Walter, Dagmar Wieczorek, Sylke Winkler, John Ziebuhr, Simone Scheithauer, Hajo Grundmann, Jonathan Schmid-Burgk, Ulrike Protzer and Inti Alberto De La Rosa Velázquez.

## Notes

### Competing Interest Statement

The authors have declared no competing interest.

