## Supplementary material for "SARS-CoV-2 infection paralyzes cytotoxic and metabolic functions of the immune cells": Suppl Info Fig

[1]Institute of Medical Genetics and Applied Genomics, University of Tübingen, Calwerstrasse 7, 72076, Tübingen, Germany
[2]NGS Competence Center Tübingen (NCCT), University of Tübingen, Calwerstrasse 7, 72076 Tübingen, Germany
[3]Werner Siemens Imaging Center, University of Tübingen, Röntgenweg 13, 72076, Tübingen, Germany
[4]Institute of Tropical Medicine, University Hospital of Tübingen, Tübingen, Wilhelmstrasse 27, 72076, Tübingen, Germany
[5]Department of Internal Medicine, Division of Endocrinology, Diabetology and Nephrology, University Hospital of Tübingen, Germany
[6]Research Institute of Women's Health, University of Tübingen, Calwerstrasse 7/6, 72076, Tübingen, Germany
*Equal contributions
### Correspondence
Dr Yogesh Singh or Prof Olaf Riess
Institute of Medical Genetics and Applied Genomics, Tübingen University
Calwerstraße 7, 72076, Tübingen, Germany


**Abstract:**

The SARS-CoV-2 virus is the causative agent of the global COVID-19 infectious disease outbreak, which can lead to acute respiratory distress syndrome (ARDS). However, it is still unclear how the virus interferes with immune cell and metabolic functions in the human body. In this study, we investigated the immune response in acute or convalescent COVID19 patients. We characterized the peripheral blood mononuclear cells (PBMCs) using flow cytometry and found that $CD8^+$ T cells were significantly subsided in moderate COVID-19 and convalescent patients. Furthermore, characterization of $CD8^+$ T cells suggested that patients with a mild and moderate course of the COVID-19 disease and convalescent patients have significantly diminished expression of both perforin and granzyme A in $CD8^+$ T cells. Using $^1$H-NMR spectroscopy, we characterized the metabolic status of their autologous PBMCs. We found that fructose, lactate and taurine levels were elevated in infected (mild and moderate) patients compared with control and convalescent patients. Glucose, glutamate, formate and acetate levels were attenuated in COVID-19 (mild and moderate) patients. In summary, our report suggests that SARS-CoV-2 infection leads to disrupted $CD8^+$ T cytotoxic functions and changes the overall metabolic functions of immune cells.

**Key words**: COVID-19, $CD8^+$ T cells, Granzyme A, Perforin, Metabolites, $^1$H-NMR, Flow cytometry

**Short title**: Defective immune-metabolic functions in COVID-19 patient

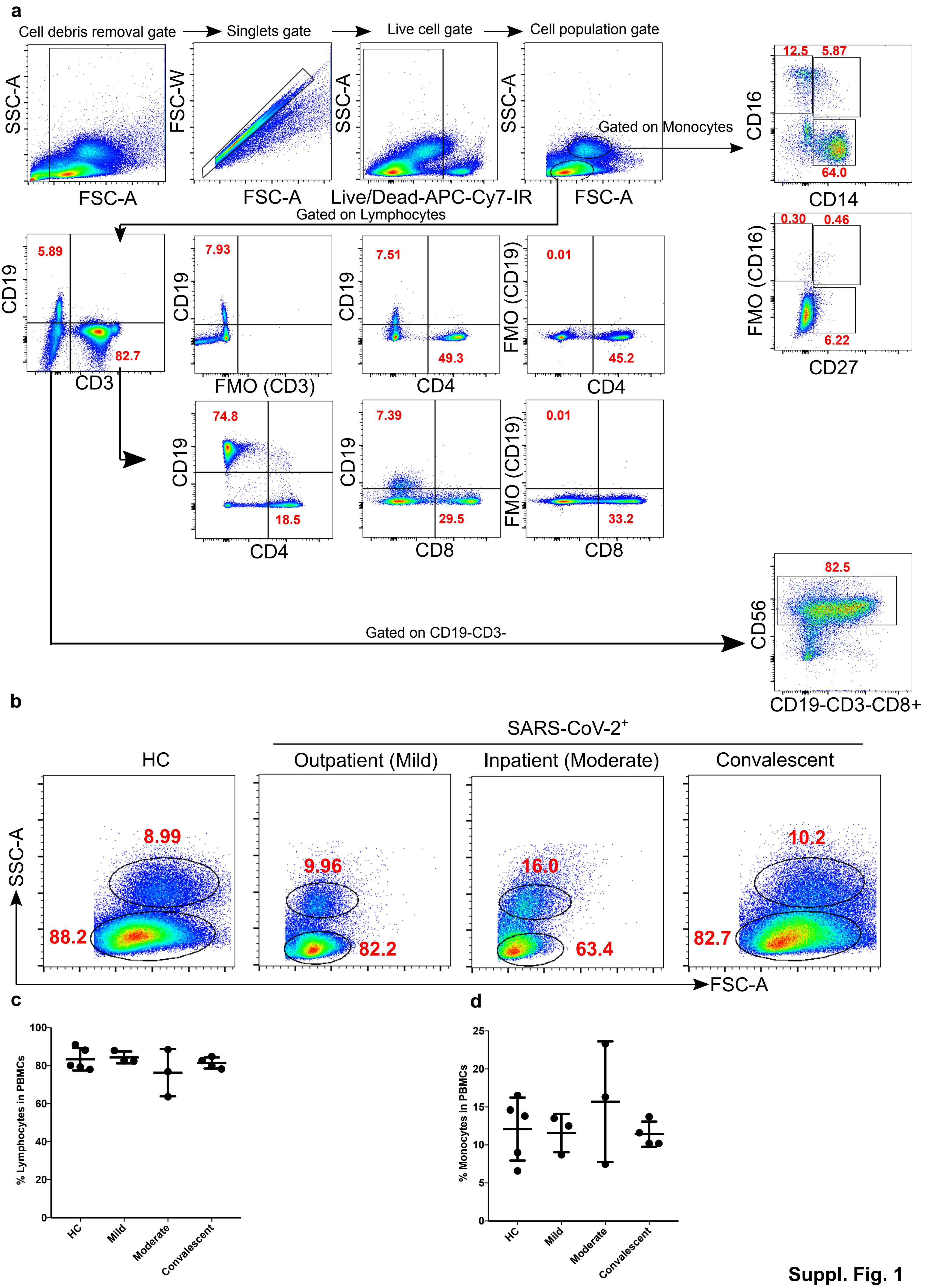
a
Cell debris removal gate
Singlets gate
Live cell gate
Cell population gate
SSC-A
FSC-A
FSC-W
Live/Dead-APC-Cy7-IR
Gated on Monocytes
Gated on Lymphocytes
CD16
CD14
12.5
5.87
64.0
FMO (CD16)
CD27
0.30
0.46
6.22
CD19
CD3
5.89
82.7
FMO (CD3)
7.93
CD4
7.51
49.3
FMO (CD19)
0.01
45.2
74.8
18.5
CD8
7.39
29.5
33.2
Gated on CD19-CD3-
CD56
CD19-CD3-CD8+
82.5
b
SARS-CoV-2+
HC
Outpatient (Mild)
Inpatient (Moderate)
Convalescent
8.99
88.2
9.96
82.2
16.0
63.4
10.2
82.7
c
% Lymphocytes in PBMCs
HC
Mild
Moderate
Convalescent
d
% Monocytes in PBMCs

**Suppl. Fig. 1**

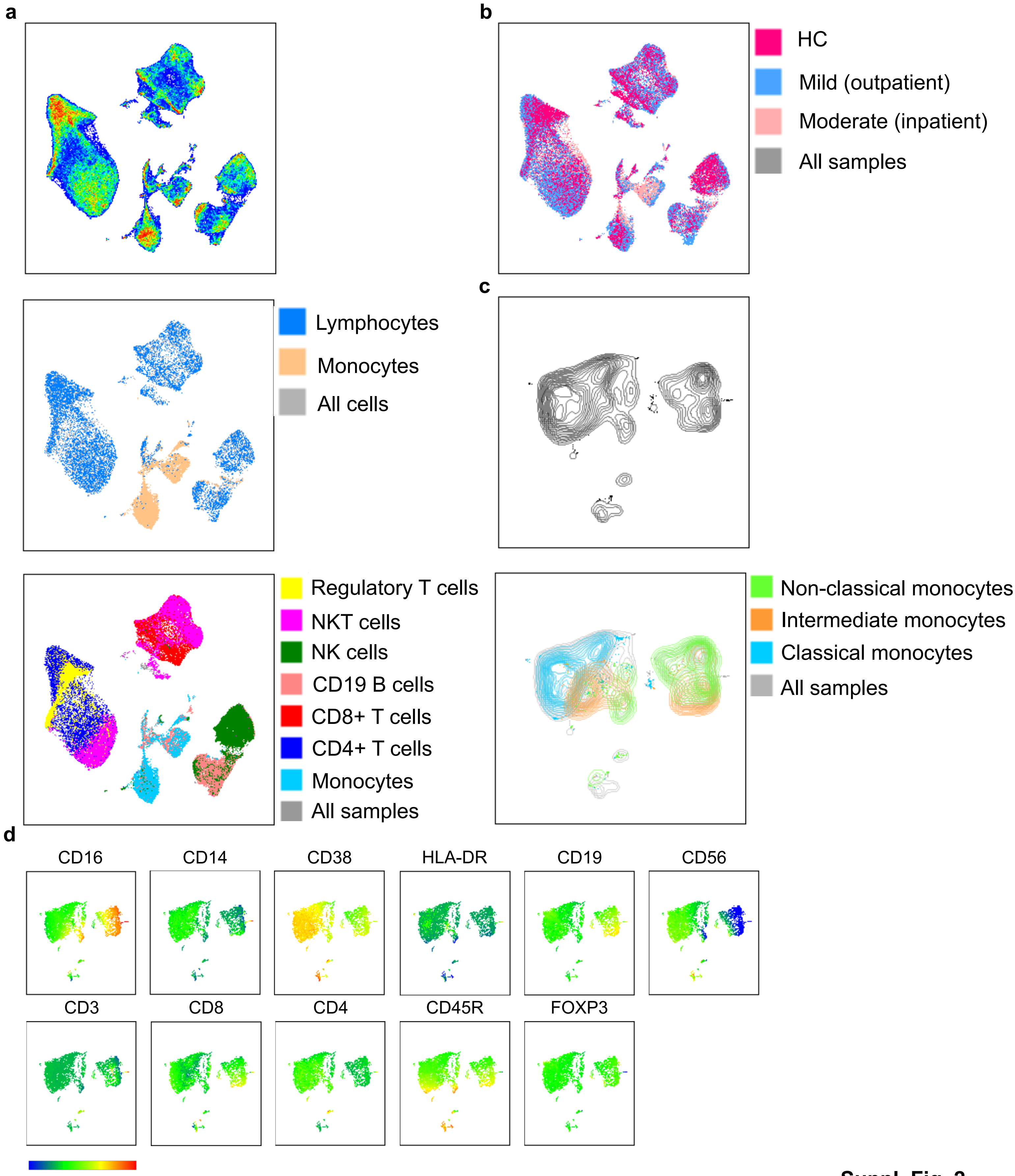
a
b
HC
Mild (outpatient)
Moderate (inpatient)
All samples
Lymphocytes
Monocytes
All cells
c
Regulatory T cells
NKT cells
NK cells
CD19 B cells
CD8+ T cells
CD4+ T cells
Monocytes
All samples
Non-classical monocytes
Intermediate monocytes
Classical monocytes
All samples
d
CD16
CD14
CD38
HLA-DR
CD19
CD56
CD3
CD8
CD4
CD45R
FOXP3

**Suppl. Fig. 2**

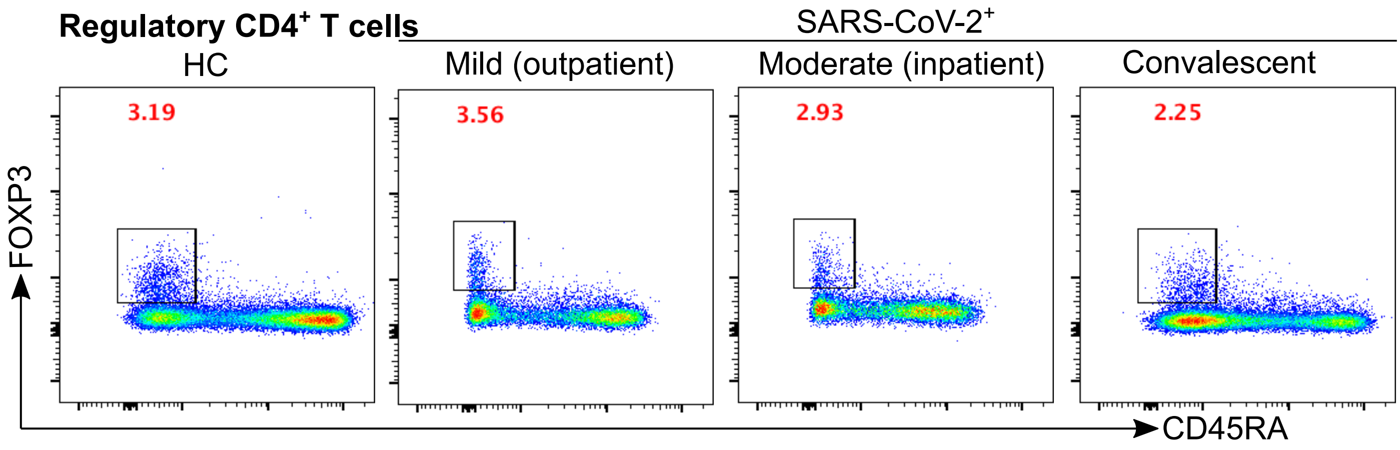
Regulatory CD4+ T cells
SARS-CoV-2+
HC
Mild (outpatient)
Moderate (inpatient)
Convalescent
3.19
3.56
2.93
2.25
FOXP3
CD45RA

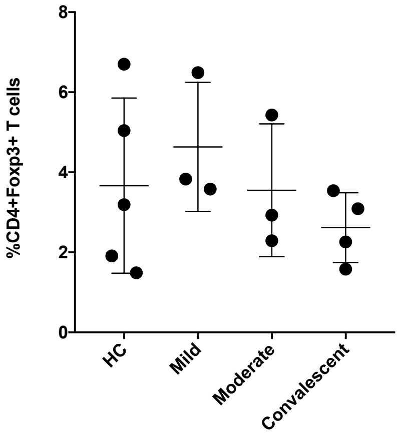
%CD4+Foxp3+ T cells
8
6
4
2
0
HC
Mild
Moderate
Convalescent

Suppl. Fig. 3

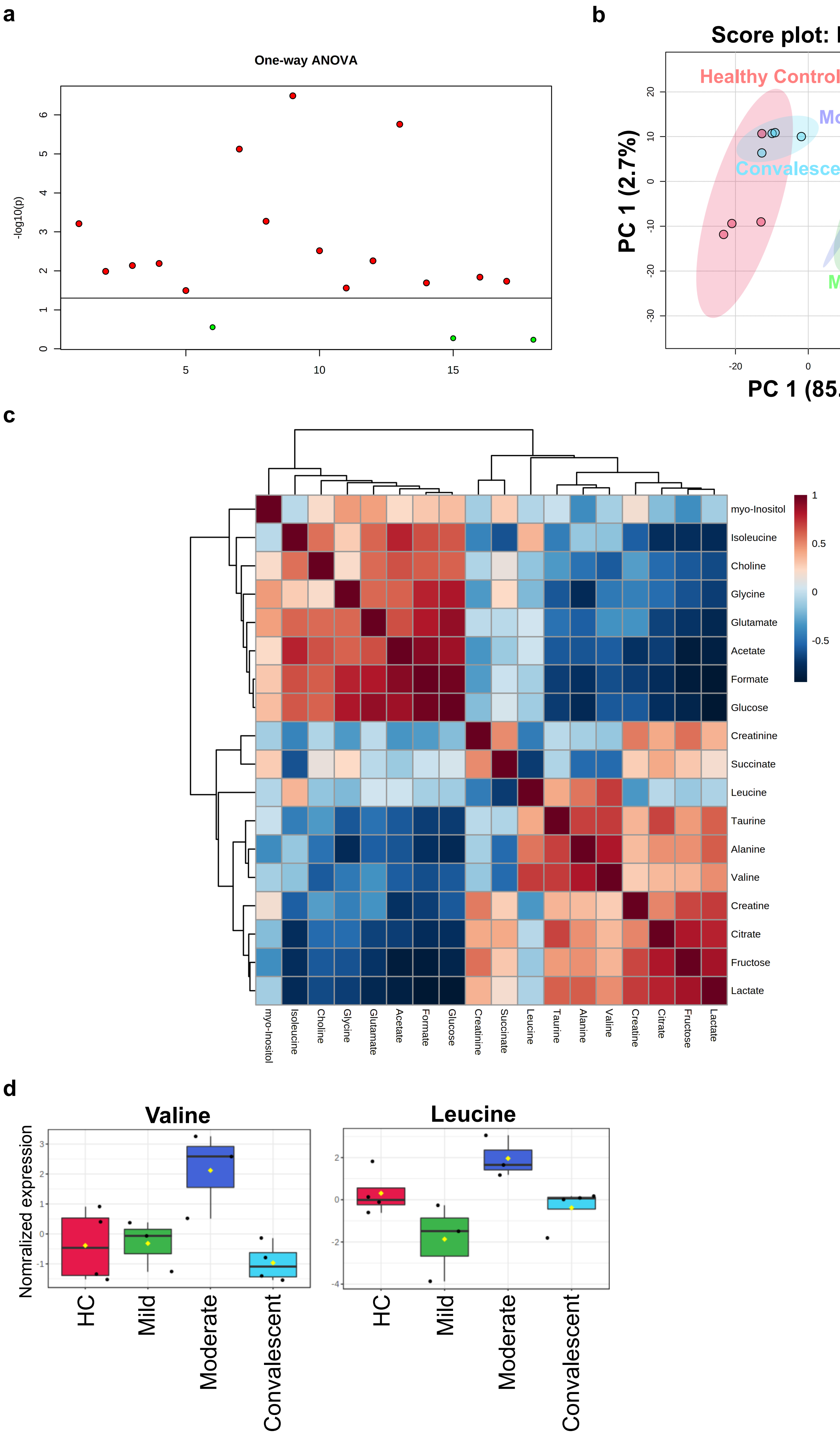
a
One-way ANOVA
-log10(p)
b
Score plot: PLSDA
1) Healthy Control
2) Mild (Outpatient)
3) Moderate (Inpatient)
4) Convalescent
Healthy Control
Moderate
Convalescent
Mild
PC 1 (2.7%)
PC 1 (85.2%)
c
myo-Inositol
Isoleucine
Choline
Glycine
Glutamate
Acetate
Formate
Glucose
Creatinine
Succinate
Leucine
Taurine
Alanine
Valine
Creatine
Citrate
Fructose
Lactate
d
Valine
Leucine
Nomralized expression
HC
Mild
Moderate
Convalescent

Suppl. Fig. 4
